# PROBind: A Web Server for Prediction, Analysis and Visualization of Protein-Protein and Protein-Nucleic Acid Binding Residues

**DOI:** 10.1101/2025.02.08.637237

**Authors:** Chaojin Wu, Fuhao Zhang, Pengzhen Jia, Jiuxiang Zhu, Min Zeng, Gang Hu, Kui Wang, Lukasz Kurgan, Min Li

## Abstract

Protein–protein and protein–nucleic acids interactions are fundamental to numerous cellular functions, yet only a small fraction have been experimentally characterized. Although modern computational methods have been developed for predicting interacting residues in proteins, they are challenging to use due to individual installation and execution requirements, lack of a standardized input or output format, and absence of support for result analysis. Moreover, methods trained using structures of complexes or intrinsically disordered regions, may not perform well on other types. To overcome these challenges, we develop PROBind, a web server for predicting, analyzing, and interactively visualizing protein, DNA and RNA binding residues from both protein sequences and structures. PROBind integrates 12 predictors trained on structural or disordered proteins, and supports the upload of results from external predictors. By normalizing and averaging predictions from multiple predictors targeting the same ligand type, PROBind generates meta-predictions that balance discrepancies among different methods. Furthermore, it provides interactive graphical tools for result analysis and contextualization. Overall, PROBind accommodates diverse ligand types and supports predictions and analysis based on both structure and sequence data, overcoming the limitations of existing tools. PROBind is freely accessible at https://www.csuligroup.com/PROBind.

## Introduction

Proteins that interact with nucleic acids and proteins contribute to many key cellular functions, such as translation, gene expression regulation, DNA transcription, and protein synthesis [1-3]. Knowledge of binding residues in proteins aids in deciphering protein functions and molecular-level mechanisms underlying diseases, and drug discovery activities [4-6]. However, the throughput of experimental methods that are used to annotate these interactions at the amino acid level does not scale with the rapidly expanding databases of protein sequences, highlighting the need for computational methods that predict interacting residues. Recent surveys reveal that numerous computational methods of protein-binding residues (PBRs), DNA-binding residues (DBRs) and RNA-binding residues (RBRs) from protein sequences and structures are available [7-14].

Current predictors can be broadly classified into two categories: structure-trained and disorder-trained. Structure-trained predictors are designed using annotations obtained from the structural information of protein–protein and protein–nucleic acid complexes. They are able to be further divided into those that make predictions using protein structures and those relying on sequences. Examples of recently developed structure-trained predictors from sequences include DeepPPISP [15], a deep learning tool that combines local (sequence window-based) and global (whole sequence-based) features to predict PBRs, and NCBRPred [16], which applies bidirectional deep LSTM-based network and multi-label learning framework to predict DBRs and RBRs. Recent structure-trained predictors from protein structures include ScanNet [17], an end-to-end geometric deep network that relies on spatio-chemical arrangement of adjacent residues and atoms to predict PBRs, and GraphBind [18], a deep graph neural network that identifies putative RBRs and DBRs. On the other hand, disorder-trained predictors focus on the intrinsically disordered regions (IDRs) in proteins – sequence regions that lack stable tertiary structures under physiological conditions [19, 20]. Consequently, these methods make predictions solely from the protein sequences. A couple of recently released representative methods in this group are a multi-task-based deep convolutional network-based DeepDISOBind [21] and a deep feedforward network-based flDPnn [22], both of which predict disordered PBRs, DBRs and RBRs. Empirical studies have identified that disorder-trained methods struggle to achieve accuracy on structure-annotated proteins, while structure-trained methods also perform poorly on intrinsically disordered proteins [23-26]. This highlights the need to integrate both types of predictors for obtaining accurate predictions.

We investigated over 50 predictors and tools of PBRs, RBRs and DBRs, focusing on practical aspects related to their availability, coverage for a broad range of scenarios (interactions with proteins, DNA and RNA), and ability to support analysis and interpretation of the resulting predictions. However, most tools fail to cover PRB, RBR, and DBR simultaneously. They usually generate predictions through a single algorithm, but studies have shown that most predictors struggle to achieve optimal performance across diverse proteins [23-26]. Furthermore, our analysis reveals that many tools provide predictions without visualization and ability for a downstream analysis, and output results in various formats, making it rather difficult to use and compare results from different tools.

In light of these observations, we developed PROBind, a web server for the prediction, visualization, and analysis of putative PBRs, RBRs and DBRs in proteins. PROBind integrates 12 predictors trained on intrinsically disordered proteins or structural proteins, and supports FASTA-formatted protein sequences or PDB-formatted protein structures as input [15-18, 21-23, 25, 27-30]. We empirically test performance of these tools on proteins with structural annotations and disordered annotations to justify inclusion of this large collection of methods. Moreover, motivated by evidence that meta-prediction can enhance predictive performance [24, 31], we examined the effect of combining outputs from multiple tools targeting the same interaction type. PROBind focuses on convenience and provides tools to analyze and contextualize predictions, allowing users to select multiple predictors, including ability to load predictions from methods that are not included in our platform, automating the entire prediction process, providing graphical panels to visualize and interact with the results, and producing additional annotations that provide useful context for the predictions.

## Results

### Overview of PROBind

PROBind is an interactive web server with a front-end built using the Vue framework (https://vuejs.org/) and a back-end developed with the SpringBoot framework (https://spring.io/projects/spring-boot). **Figure 1** summarizes the workflow of PROBind including input formats, predictive methods employed, downstream analysis of predictions and result visualisation. PROBind accepts input either as FASTA-formatted protein sequences or PDB-formatted structures, and the web server correspondingly highlights for selecting the predictors that utilize these inputs. Users may paste their input data into a text field or upload a file containing the data. The web server supports the submission of multiple sequences or a single protein structure at a time; the lower limit for the structure input is due to a longer runtime of methods that predict from structures. The input page provides examples and hints that explain format and other details of the inputs.

**Figure 1.**
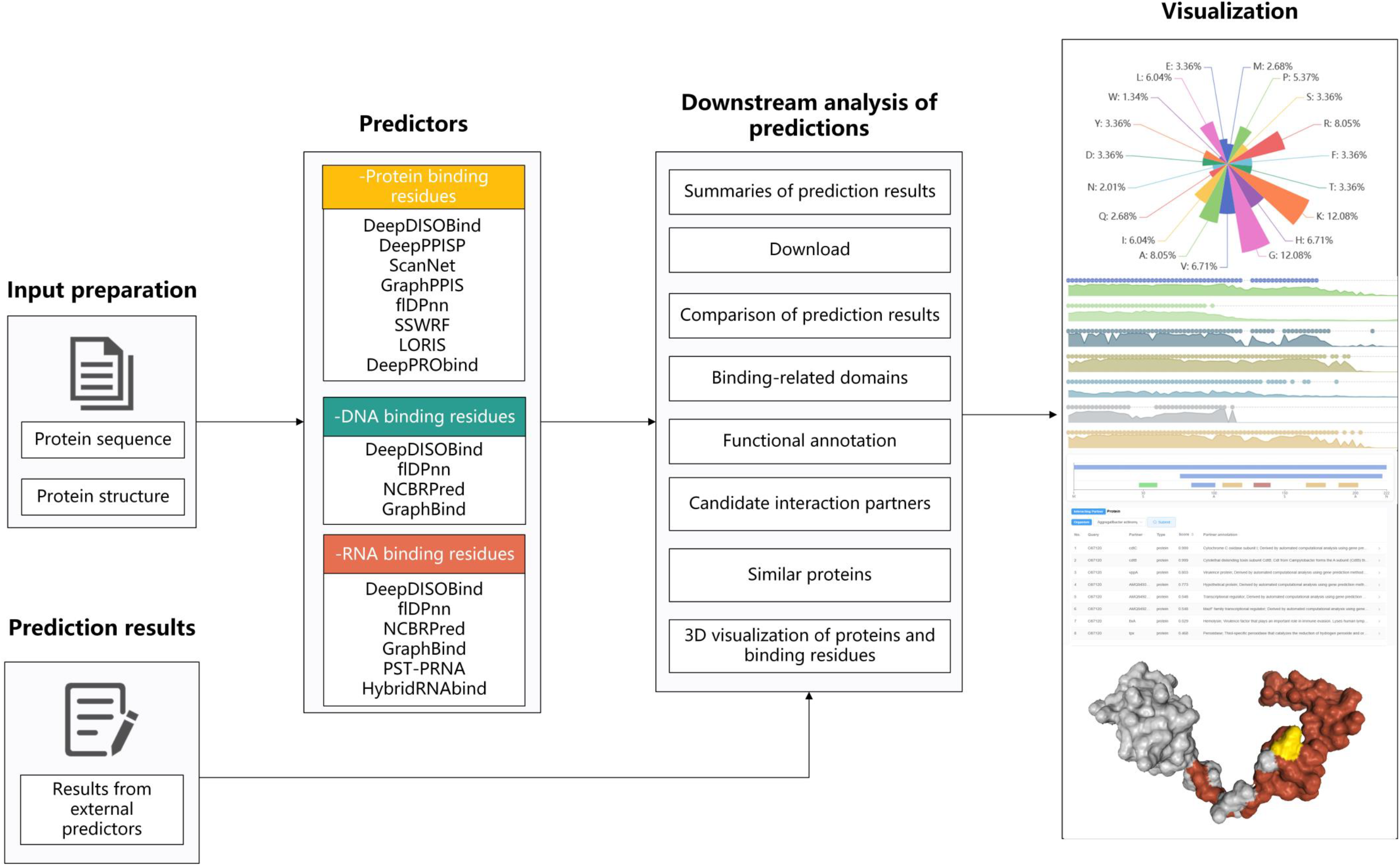
Workflow of the PROBind web server.

PROBind generates predictions from 12 integrated methods, allowing users to choose any subset of these predictors. Given the relatively high computational cost of the included structure-trained methods, we do not support predictions where sequences are used to predict structure with AlphaFold [32] to subsequently use methods that predict binding residues from structure. However, users can produce and utilize the AlphaFold-generated structure as the input. To facilitate selection of appropriate methods, PROBind suggests predictors that can be used when the input is sequence or structure. The sequence-based predictors can be still used when the structure is the input. To reduce the runtime, PRObind runs the selected predictors in parallel and updates the status of the prediction process in real time. After the predictions are finalized, the server automatically redirects to a page that offers a comprehensive downstream analysis of the results. If users provide an email address on the input page, the server will send a notification email to inform users when the predictions are ready, specifying their location, and including text files containing all prediction results generated by the selected predictors.

### Downstream analysis page

While predictors originally output results using different formats, the downstream analysis page unifies these formats and provides one combined visualization that includes residue-level and aggregate/protein-level results. PROBind also supports uploading results from external predictors. We explain details of format of the files that can be uploaded in **Suppl. File S1**. The result page includes a menu on the left that can be used to find a specific part of the results and to adjust thresholds used to derive binary predictions of binding residues. The top of the result page summarizes protein-level results (e.g. overall fraction of putative binding residues) for each ligand type (DNA, RNA and protein) and each selected predictor. It also provides access to text files with the predictor-generated results. Below it shows residue-level predictions by interactive graphical plots. For instance, using these graphical plots users can zoom on specific parts of the sequence, read details of predictions for specific residues (predicted propensities and binary predictions of PBRs, DBRs and RBRs) by hovering over them. The results page also includes a meta-prediction that averages results from methods that address prediction of the same ligand type to ease and improve identification of putative binding residues. Users can select which tools are used for the meta-prediction.

The bottom of the result page provides relevant context for the predictions. It visualizes annotations generated with InterProScan [33, 34] and ProtENN [35]. InterProScan produces functional regions including protein domains and motifs. We show these annotations using an interactive color-coded graph where users can hover over specific domains/motifs to see their details including location in the sequence, associated GO terms [36, 37], and their types, accession and description from InterPro. ProtENN predicts Pfam terms [38] using multiple ProtCNN models. We analyze the top five terms from each model and produce the term with the highest frequency as predicted function. We also list GO terms that correspond to the predicted Pfam terms. Next, the page provides interacting partner molecules that we obtain from the STRING [39] and BioLiP [40] databases. The server uses Diamond[41] to identify the most similar protein in interaction database based on the input protein. **Suppl. File S2** shows the details how PROBind generates the interacting partners. We show the partner data using a tabular format that includes identifier and type of partner molecule, score that quantifies reliability of this annotation, and a brief description of the interaction. Finally, the page ends with the list of similar proteins.

The results page also generates 3D model of protein structure with color-coded annotations of putative binding residues if structure is used as the input. We use NGL Viewer [42] which provides a user-friendly interface that allows users to adjust how structure is visualized, including the colour scheme to annotate putative binding residues. The structure view is supplemented with a table that provides predicted propensities and binary predictions for all amino acids.

### Empirical analysis of predictive performance

We utilize three test datasets containing PBRs, RBRs and DBRs with structural annotations and disordered annotations to evaluate the predictive performance of 12 methods included in PROBind. We also provide a meta-predictor, which averages normalized (to a unit range) propensities generated by the user-selected predictors. User can choose any subset of methods that they originally selected to generate predictions. Here, we consider two meta-predictors: *MetaAll* that averages propensities generated by all methods that predict a given type of interaction (PBRs, RBR, and DBRs), and *MetaSelected* that uses a subset of newer and more accurate predictors that include structure- and disorder-trained tools. **Suppl. Table S1** shows the predictors that we use to implement the two meta-methods. To quantify predictive performance, we use the area under the receiver operating characteristic curve (AUC), a widely metric used in related studies. We summarize results for the 12 predictors and 2 meta-predictors in **Figure 2**.

**Figure 2.**
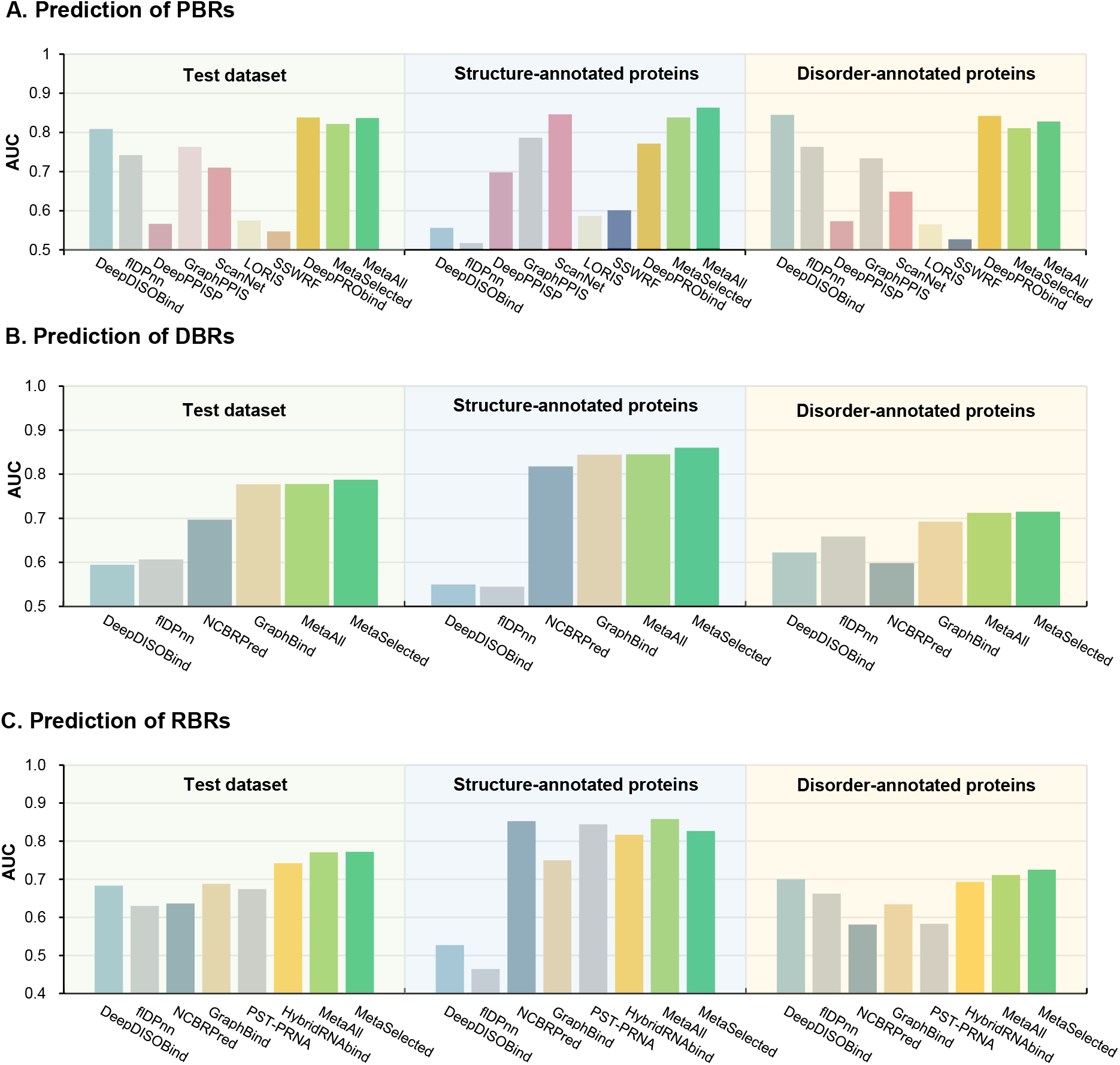
Predictive performance of the 12 predictors. Predictive performance of the 12 predictors and 2 selected meta-predictors on the test datasets and their subsets that include the structure-annotated and the disorder-annotated proteins. **Panel A**: Comparative assessment of predictors on proteins-binding residues (PBRs) test dataset. **Panel B**: Comparative assessment of predictors on DNA-binding residues (DBRs) test dataset. **Panel C**: Comparative assessment of predictors on RNA-binding residues (RBRs) test dataset.

We find that current predictors struggle to produce equally accurate results for different proteins. For instance, in the prediction of PBRs (**Figure 2A**), DeepDISOBind achieves the best performance on proteins with disordered annotations (AUC = 0.845), but drops to 0.556 on proteins with structural annotations. By contrast, the best structure-trained predictor, ScanNet, achieves an AUC of 0.846 for the structure-annotated proteins but only 0.649 for disorder-annotated proteins. Notably, DeepPRObind demonstrates relatively minor performance discrepancies between these two protein categories, likely due to its unique training strategy, which employs distinct datasets and training modules for different proteins. A similar trend is observed when evaluating RNA-binding residue predictors (**Figure 2C**). The structure-trained predictor NCBRPred achieves the top AUC of 0.853 on the proteins with structural annotations but plummets to 0.581 on the proteins with disordered annotations. Conversely, the top-performing disorder-trained method, DeepDISOBind, achieves an AUC of 0.700 on disordered proteins yet only achieves AUC=0.527 on the structural proteins. HybridRNAbind is a meta-predictor based on DeepDISOBind and NCBRPred, where DeepDISOBind specializes in predicting RNA binding residues within disordered proteins, while NCBRPred targets in structural proteins. By integrating outputs from both tools, HybridRNAbind not only achieves higher overall predictive power but also demonstrates top performance on the entire test set with minimal performance gaps between subsets. An exception arises in DNA-binding residue predictions **(Figure 2B**), where the structure-trained predictor, GraphBind, exhibits the highest AUC of 0.844 and 0.692 for the structural proteins and disordered proteins with structural annotations and disordered annotations, respectively. However, NCBRPred, also trained on structural proteins, secures an AUC of 0.818 for structural proteins, but obtains the lowest AUC of 0.598 on the disordered proteins. Overall, these results show that while some predictors provide relatively accurate predictions, methods trained on disordered proteins generally underperform on structural proteins, and methods trained on structural proteins yield lower-quality predictions for disordered proteins — consistent with prior observations [23-26]. Consequently, combining multiple methods, particularly those trained on both structural and disordered proteins, may yield more robust predictions across different protein types. This insight motivates our development and evaluation of meta-predictors in PROBind.

The meta-predictors (green bars in **Figure 2**) show better balance across both structural and disordered proteins compared to 12 individual tools. For predicting protein-binding residues (**Figure 2A**), *MetaAll* secures AUCs of 0.838 and 0.811 for structural and disordered proteins, respectively. When predicting DNA-binding residues (**Figure 2B**), *MetaAll* achieves AUCs of 0.845 and 0.712, and for RNA-binding residues (**Figure 2C**), it achieves AUCs of 0.858 and 0.711. We find a similar pattern for the *MetaSelected* meta-predictor, whose results closely align with the top-performing structure-trained predictor on the structural subset and the top disorder-trained predictor on the disordered subset. When evaluated on the complete dataset, *MetaAll* produces AUCs of 0.822, 0.778 and 0.771 for PBRs, DBRs and RBRs, respectively, compared to the best results from the individual methods that are 0.838 for PBRs (DeepPRObind), 0.777 for DBRs (GraphBind) and 0.742 for RBRs (HybridRNAbind). This shows that the more-balanced predictions of the meta-methods match the quality of the best predictors used individually. The bottom line is that we recommend that the users should generate the meta-prediction that includes one or (preferably) more structure-trained and disorder-trained methods if they do not know whether their target proteins are disordered or structural.

Furthermore, we found that disorder-trained predictors are less than structure-trained predictors, particularly in PBR predictors. The PBR predictors contains 5 structure-trained methods (DeepPPISP, SSWRF, LORIS, ScanNet, and GraphPPIS), 2 disorder-trained methods (flDPnn, and DeepDISOBind) and 1 method trained on both structural and disordered proteins (DeepPRObind). To investigate the impact of this imbalance on fusion-based prediction, we conducted additional analyses, summarized in **Tables 1-3**. For PBR predictors, DeepPRObind, DeepDISOBind, and ScanNet demonstrated the highest performance on the overall test dataset, structural dataset, and disordered dataset, respectively. MetaSelected combines the results from these three top-performing predictors, while MetaAll combines the results from all methods of PBRs. We observed that MetaSelected achieved comparable results to MetaAll, and even outperformed MetaAll in the prediction of PBRs and DBRs. This suggests that integrating appropriate methods for meta-prediction is more useful than simply combining a larger number of methods to improve prediction accuracy. Moreover, similar trends were observed in the meta-predictions of DBRs and RBRs, as shown in **Tables 2 and 3**. Therefore, we recommended the best-performing methods for different protein types and binding partner types, rather than integrating all methods.

**Table 1.** Meta-predictions based on diverse predictors of protein-binding residues.

| Methods | Test dataset | Structure-annotated test proteins | Disorder-annotated test proteins |
| --- | --- | --- | --- |
| DeepPRObind | 0.838 | 0.771 | 0.842 |
| DeepDISOBind | 0.809 | 0.556 | 0.845 |
| ScanNet | 0.710 | 0.846 | 0.649 |
| DeepDISOBind+ScanNet | 0.808 | 0.848 | 0.794 |
| MetaSelected | 0.837 | 0.863 | 0.828 |
| MetaAll | 0.822 | 0.838 | 0.811 |

*Note:* 1. DeepPRObind, DeepDISOBind, and ScanNet: the top-performing predictor on the entire test dataset, structure-annotated test proteins and disorder-annotated test proteins, respectively; 2. MetaSelected combines results generated by DeepPRObind, DeepDISOBind, and ScanNet; 3. MetaAll combines results generated by all predictors.

**Table 2.**
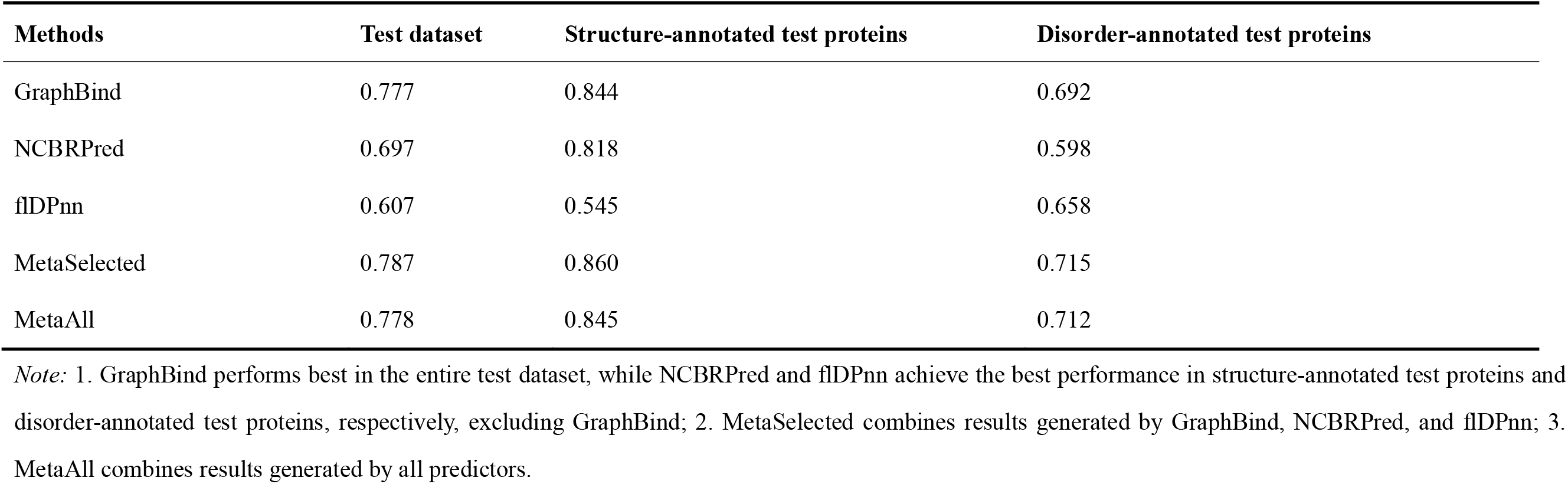
Meta-predictions based on diverse predictors of DNA-binding residues.

**Table 3.** Meta-predictions based on diverse predictors of RNA-binding residues.

| Methods | Test dataset | Structure-annotated test proteins | Disorder-annotated test proteins |
| --- | --- | --- | --- |
| HybridRNAbind | 0.742 | 0.817 | 0.693 |
| NCBRPred | 0.636 | 0.853 | 0.581 |
| DeepDISOBind | 0.683 | 0.527 | 0.700 |
| MetaSelected | 0.772 | 0.827 | 0.725 |
| MetaAll | 0.771 | 0.858 | 0.711 |

*Note:* 1. HybridRNAbind, NCBRPred, and DeepDISOBind: the top-performing predictor in the entire test dataset, structure-annotated test proteins and disorder-annotated test proteins, respectively; 2. MetaSelected combines results generated by DeepDISOBind, flDPnn, NCBRPred, GraphBind, and HybridRNAbind; 3. MetaAll combines results generated by all predictors.

### Case study

We use Cytolethal distending toxin subunit A (CDTA, UniProt accession: O87120, DisProt: DP01002) as a case study to describe the visualization capabilities of PROBind web server. This protein is used as one of the example inputs on the server page. We use protein structure as the input and select all predictors of PBRs (input page shown in **Suppl. Figure S1**) since the protein is known to interact with Cytolethal distending toxin nuclease subunit Aa-CdtB and Cytolethal distending toxin subunit Aa-CdtC [39, 43]. **Suppl. Figures S2-S5** show the different types of downstream analysis. **Suppl. Figure S2** shows an overview of the prediction at the protein level, which indicates that this protein contains protein-binding regions from 13 to 146. **Suppl. Figure S3** shows the prediction results of eight methods for PBRs (DeepPPISP, GraphPPIS, ScanNet, SSWRF, LORIS, DeepPRObind, DeepDISOBind and flDPnn). **Suppl. Figure S3** reveals that the first 70 amino acids at the start of the sequence are bound to proteins, and this coincides with the annotation information of DisProt. This is further supported by the MetaAll and MetaSelected predictions shown in **Figure 3**.

**Figure 3.**
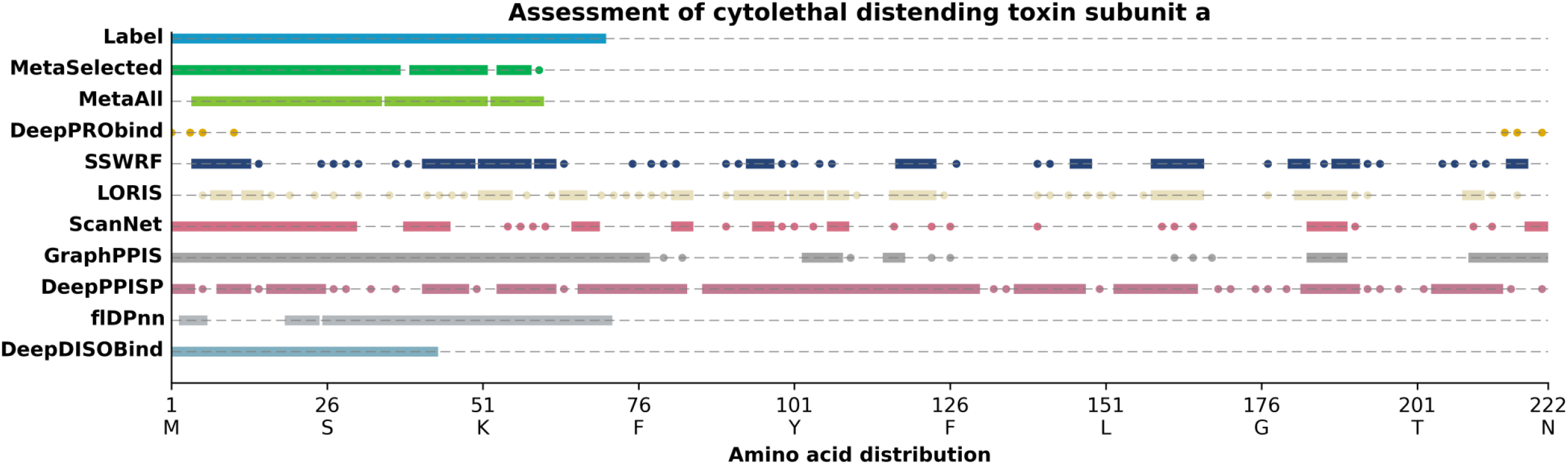
Case study of Cytolethal distending toxin subunit A (CDTA, UniProt accession: O87120, DisProt: DP01002)

To provide a useful context, the downstream analysis page offers functional annotations and a list of interacting partners extracted from other resources, as we detail above. These functional annotations show that the K1 to G70 region is annotated with three GO terms: GO:0005179, and GO:0005576 (**Suppl. Figure S4**). **Suppl. Figure S5** reveals that the external resources, i.e., STRING [39] and BioLip [40] databases, identify that this protein interacts with 8 partners, with notable emphasis on cdtB and cdtC. Finally, **Suppl. Figure S6** visualizes the predicted PBRs in an interactive view of the structure of this protein.

## Materials and methods

### Selection of predictors

Our study applies a two-step process to select predictors for inclusion in PROBind. First, we consider methods that can be imported into our platform. This converts into the following two criteria: 1) working open-source code; 2) results must include real-valued propensities for validation. The second step ensures that the methods: 1) cover predictions of PBRs, RBRs or DBRs; 2) are trained from structural or from intrinsic disordered annotations; and 3) use protein structure or sequence as the input. Consequently, we select 12 tools (8 structure-trained, 2 disorder-trained, and 2 structure and disorder-trained) that we summarize in **Suppl. Table S2**. There are 4 methods that use structures as input: ScanNet [17] and GraphPPIS [28] that predict PBRs, GraphBind [18] that predicts DBRs and RBRs, and PST-PRNA [29] that predicts RBRs. The other 6 tools that predict from sequences are structure-trained predictors of PBRs: DeepPPISP [15], SSWRF [27], and LORIS [30], and structure-trained NCBRPred [16] that predicts DBRs and RBRs; and disorder-trained DeepDISOBind [21] and flDPnn [22] that predict disordered PBRs, RBRs and DBRs. HybridRNAbind [23] and DeepPRObind [25] trained on both disordered and structural proteins, predict PBRs and RBRs, respectively.

### Datasets

We construct three test datasets with annotations of PBRs, DBRs and RBRs, each of which consists of proteins with structural and disordered annotations. First, we collect test datasets from DeepPRObind and HybridRNAbind, which covers disordered and structural proteins with PBRs and RBRs. For the DNA-binding residue test dataset, we combined disordered proteins sourced from the DisProt [43] database and the DeepDISOBind test dataset, and structural proteins from the GraphBind test dataset. Second, we obtain training datasets of all integrated predictors. Third, we use CD-HIT [44] to remove proteins in test datasets that share similarity > 30% with proteins in training datasets; this ensures that the remaining proteins are dissimilar to the data used to train the selected predictors. Consequently, we collect 99 proteins for the PBRs test dataset (66 disordered proteins and 33 structural proteins); 336 proteins for the DBR test dataset (168 disordered proteins and 168 structural proteins); and 409 proteins for the RBR test dataset (204 disordered proteins and 205 structural proteins). A summary of these datasets is provided in **Suppl. Table S3**.

## Data availability

PROBind and all test datasets are freely accessible at https://www.csuligroup.com/PROBind.

## CRediT author statement

**Chaojin Wu:** Software, Methodology, Visualization, Writing - original draft, Writing - review & editing, Investigation. **Fuhao Zhang:** Methodology, Writing - original draft, Writing - review&editing. **Pengzhen Jia:** Methodology, Investigation. **Jiuxiang Zhu:** Visualization. **Min Zeng:** Writing - review&editing. **Gang Hu:** Methodology. **Kui Wang:** Methodology. **Lukasz Kurgan:** Writing - review&editing, Methodology. **Min Li:** Supervision, Project administration, Funding acquisition, Writing - review & editing.

All authors have read and approved the final manuscript.

## Acknowledgements

This work was supported by the National Natural Science Foundation of China [No. 61832019 to M.L.], Hunan Provincial Science and Technology Program [2019CB1007 and 2021RC4008 to M.L.].

