## Supplementary material for "PROBind: A Web Server for Prediction, Analysis and Visualization of Protein-Protein and Protein-Nucleic Acid Binding Residues": Suppl. File S1

**Suppl. File S1. Format of the files with predictions**

To support uploading results from different predictors, we design a flexible input interface and use the following keywords to identify contents of specific rows or columns in the file: Protein Identifier, Sequence, Amino Acid, Protein Propensity, Protein Binary, DNA Propensity, DNA Binary, RNA Propensity, RNA Binary, and #. Protein Identifier is the unique name/description of the input protein. Sequence is the amino acid sequence for results in the row-wise format while Amino Acid indicates the consecutive amino acid types when the results are represented in the column-wise format. Protein binary is the binary predictions for protein binding. Protein Propensity is the predicted propensities for PBRs. Similarly, DNA Propensity, DNA Binary, RNA Propensity and RNA Binary represent predictions of DBRs and RBRs. The propensity values are comma-separated. For comments, users can use the “#” character and the webserver will skip this column or line when processing uploaded data.

We use the outputs from DisoRDPbind and NucBind as examples, see top and bottom panels in **Figure 1 below**, respectively. The top panel in **Figure 1** represents predictors, like DisoRDPbind, that output predictions in the row-wise format. The results of each protein are composed of 8 lines: the protein identifier starting with ‘>’ symbol, the amino acid sequence, binary predictions (binding vs. non-binding) for RNA-binding, comma-separated putative propensities for RNA-binding, binary predictions for DNA-binding, comma-separated putative propensities for DNA-binding, binary predictions for protein-binding, and the comma-separated putative propensities for protein-binding. We also include a descriptive title at the beginning of each line that identifies the information that is includes. In addition, PROBind accepts uploads of results in a column-wise format, like the NucBind’s predictions (bottom panel in **Figure 1 below**). The first row is the protein identifier that start with the ‘>’ symbol. The subsequent rows are predictions. The first column lists the indexes of amino acids in the input protein sequence. The second, third and fourth columns provide the consecutive amino acid types, binary predictions (binding vs. non-binding), and putative propensities for DNA-binding, respectively.

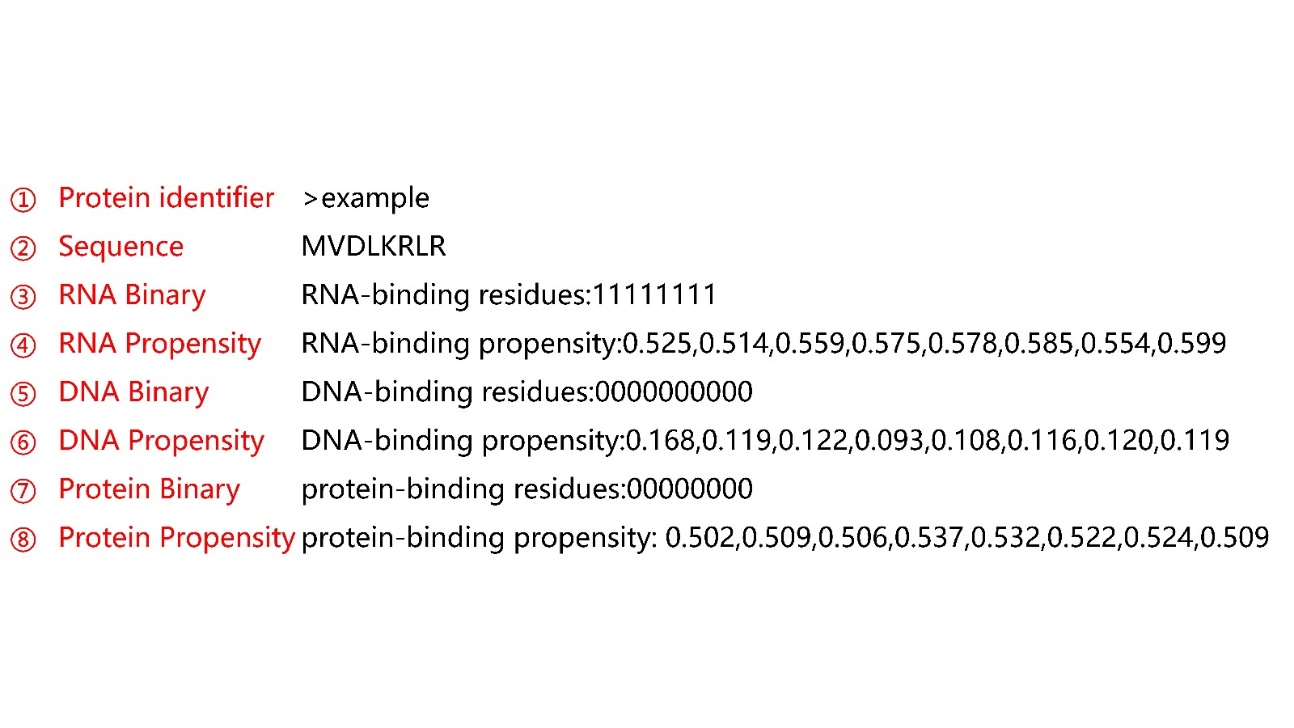

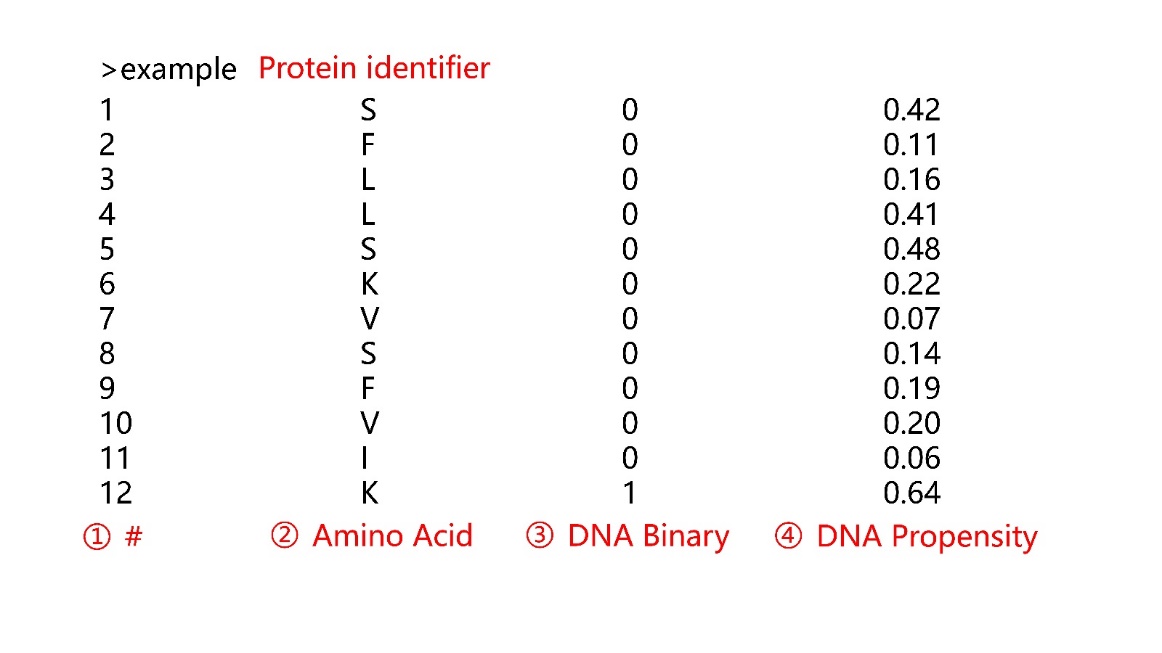

**Figure 1 Examples of uploading results from predictors outside PROBind**
