## Supplementary material for "PROBind: A Web Server for Prediction, Analysis and Visualization of Protein-Protein and Protein-Nucleic Acid Binding Residues": Suppl. File S2

**Suppl. File S2** **Generation of interacting partners**

**Figure 1 below** shows the process how PROBind generates the interacting partners. PROBind identifies protein- and nucleic acid-interacting partners based on the STRING [1] and BioLiP [2] databases, respectively. The server uses Diamond [3] to identify the most similar protein in the STRING database based on the input sequence/structure and organism, and utilizes this most similar protein to search for its interacting partners. We compute a score that estimates reliability of these interactions by multiplying the value of similarity between proteins (the user-provided input protein and the most similar protein from STRING) by the interaction score generated by STRING; similarity and interaction score range between 0 and 1, and consequently our score is in the same range. To extract the nucleic acid-interacting partners, we generate a dataset of proteins that interact with DNA/RNA in the BioLiP database and remove redundancy. Similar as for protein-interacting partners, PROBind uses Diamond to look up similar proteins in the nucleic acid-interacting database and output their interacting partners as the possible nucleic acid-interacting partners for the input protein. The web server uses the value of similarity between the user-provided input protein and the most similar protein as the score that estimates reliability of the produced protein-nucleic acid interactions.

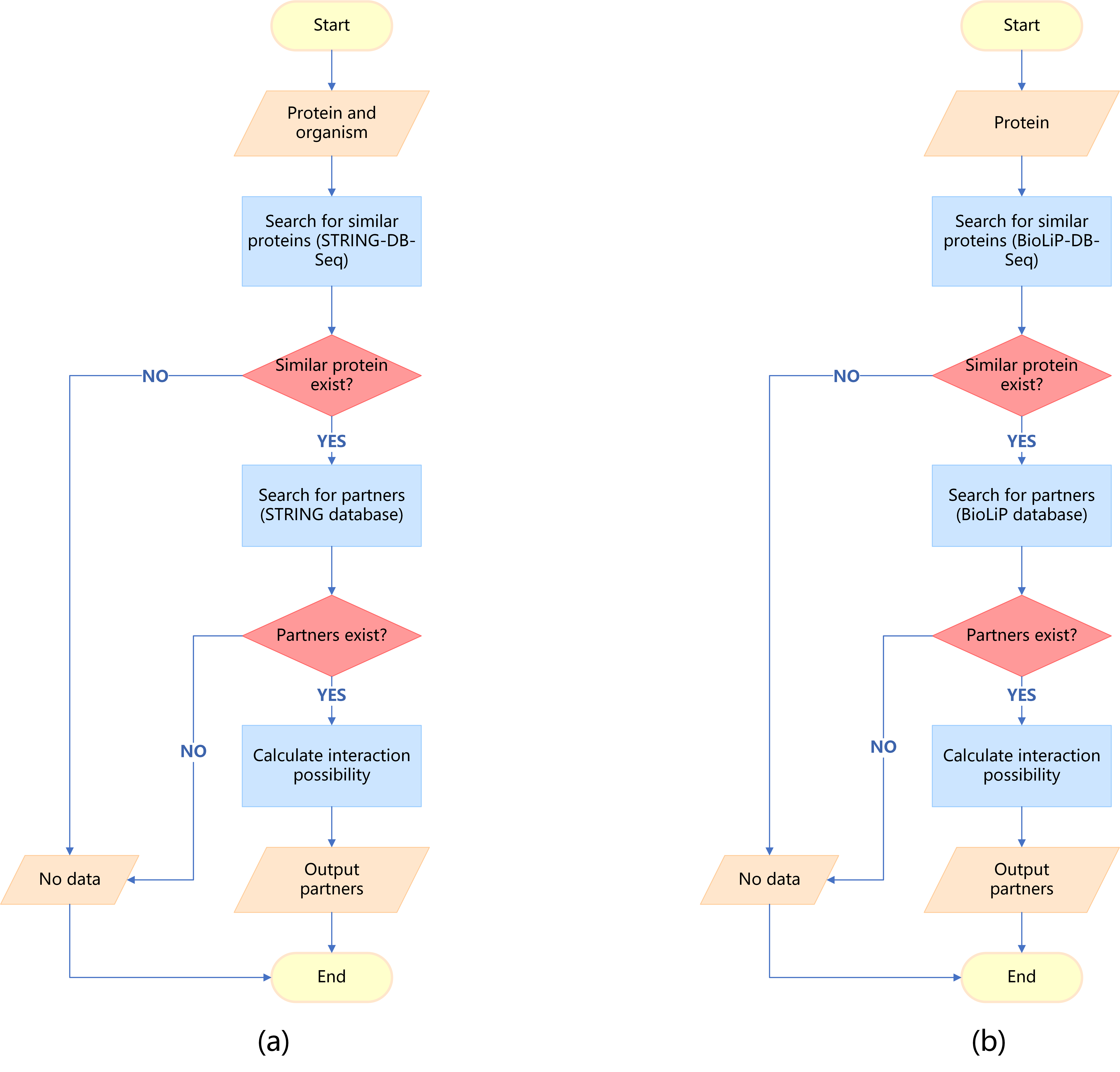

**Figure 1 Calculation of interacting partners.** (a) Protein–protein interaction partners; (b) Protein–nucleic acid interaction partners.
