## Supplementary material for "PROBind: A Web Server for Prediction, Analysis and Visualization of Protein-Protein and Protein-Nucleic Acid Binding Residues": Suppl. Table S1

**Suppl. Table S1 Predictors that PROBind uses to derive the MetaAll and MetaSelected predictors.**

| **Prediction target** | **Meta-predictors** | **Source predictors** |
| --- | --- | --- |
| Protein-binding residues | MetaAll | DeepDISOBind, DeepPPISP, GraphPPIS, flDPnn, ScanNet, LORIS, SSWRF, DeepPRObind |
|  | MetaSelected | DeepDISOBind, ScanNet, DeepPRObind |
| DNA-binding residues | MetaAll | DeepDISOBind, flDPnn, GraphBind, NCBRPred |
|  | MetaSelected | flDPnn, GraphBind, NCBRPred |
| RNA-binding residues | MetaAll | flDPnn, GraphBind, PST-PRNA, DeepDISObind, NCBRPred, HybridRNAbind |
|  | MetaSelected | flDPnn, GraphBind, DeepDISOBind, NCBRPred, HybridRNAbind |
