## Supplementary material for "PROBind: A Web Server for Prediction, Analysis and Visualization of Protein-Protein and Protein-Nucleic Acid Binding Residues": Suppl. Table S2

**Suppl. Table S2** **The predictors included in PROBind.**

| **Method** | **Input** | **Type** | **Predictive target** |
| --- | --- | --- | --- |
| ScanNet | Structure | Structure-trained | Protein-binding residues |
| GraphPPIS | Structure | Structure-trained | Protein-binding residues |
| GraphBind | Structure | Structure-trained | DNA/RNA-binding residues |
| PST-PRNA | Structure | Structure-trained | RNA-binding residues |
| DeepPPISP | Sequence | Structure-trained | Protein-binding residues |
| SSWRF | Sequence | Structure-trained | Protein-binding residues |
| LORIS | Sequence | Structure-trained | Protein-binding residues |
| NCBRPred | Sequence | Structure-trained | DNA/RNA-binding residues |
| flDPnn | Sequence | Disorder-trained | Protein/DNA/RNA-binding residues |
| DeepDISOBind | Sequence | Disorder-trained | Protein/DNA/RNA-binding residues |
| DeepPRObind | Sequence | Structure/Disorder-trained | Protein-binding residues |
| HybridRNAbind | Sequence | Structure/Disorder-trained | RNA-binding residues |
