## Supplementary material for "PROBind: A Web Server for Prediction, Analysis and Visualization of Protein-Protein and Protein-Nucleic Acid Binding Residues": Suppl. Table S3

**Suppl. Table S3 The sources and details for the three test datasets.**

| **Type** | **References used to source test datasets** | **Disorder** | | **Structure** | | **Total proteins** |
| --- | --- | --- | --- | --- | --- | --- |
|  |  | **Binding proteins** | **Non-binding proteins** | **Binding proteins** | **Non-binding proteins** |  |
| Protein | DeepPRObind | 50 | 16 | 21 | 12 | 99 |
| DNA | DeepDISOBind, DisProt, GraphBind | 21 | 147 | 86 | 82 | 336 |
| RNA | HybridRNAbind | 18 | 186 | 14 | 191 | 409 |
