## Supplementary figures and images for "PROBind: A Web Server for Prediction, Analysis and Visualization of Protein-Protein and Protein-Nucleic Acid Binding Residues"

### Suppl. Figure S1

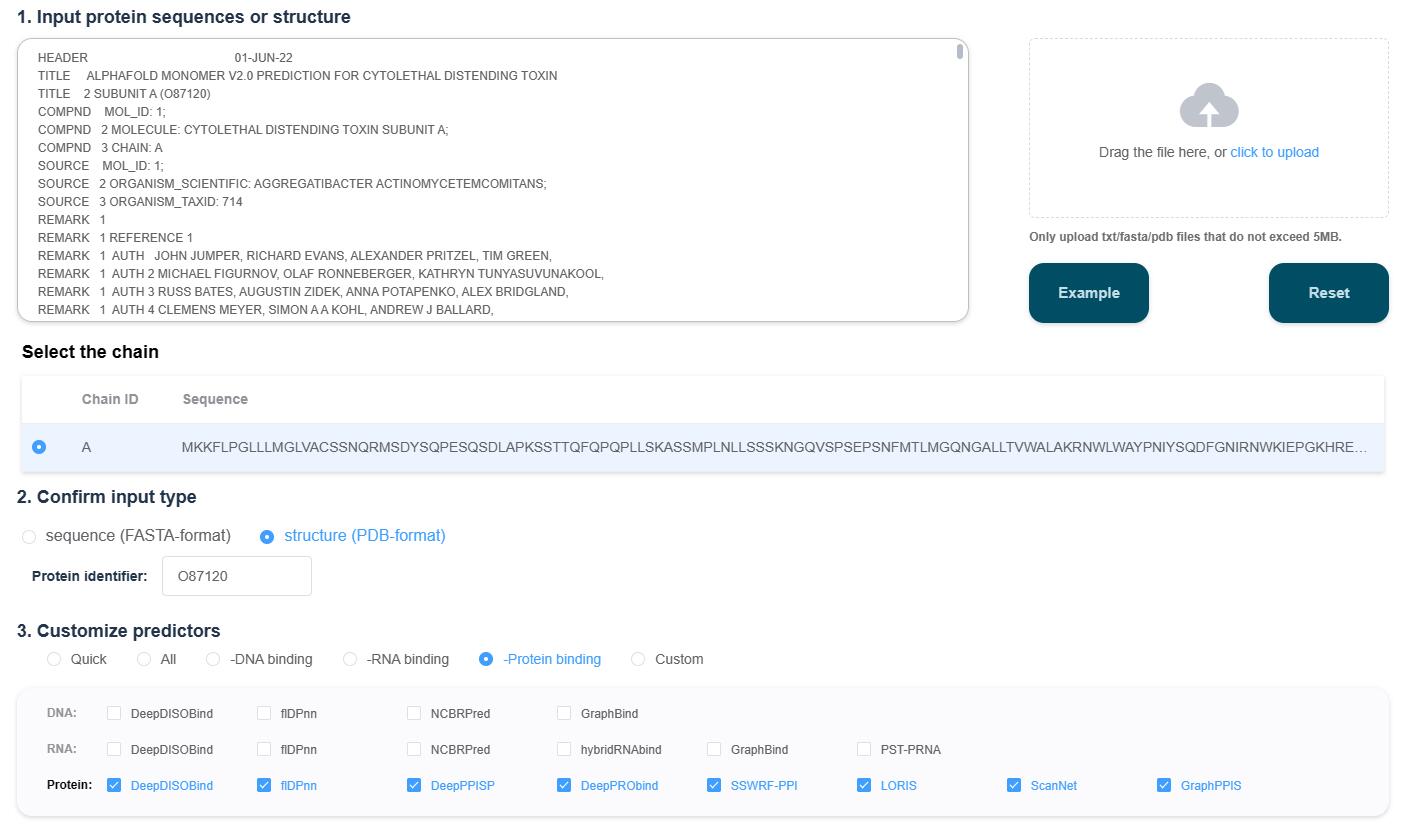

### Suppl. Figure S2

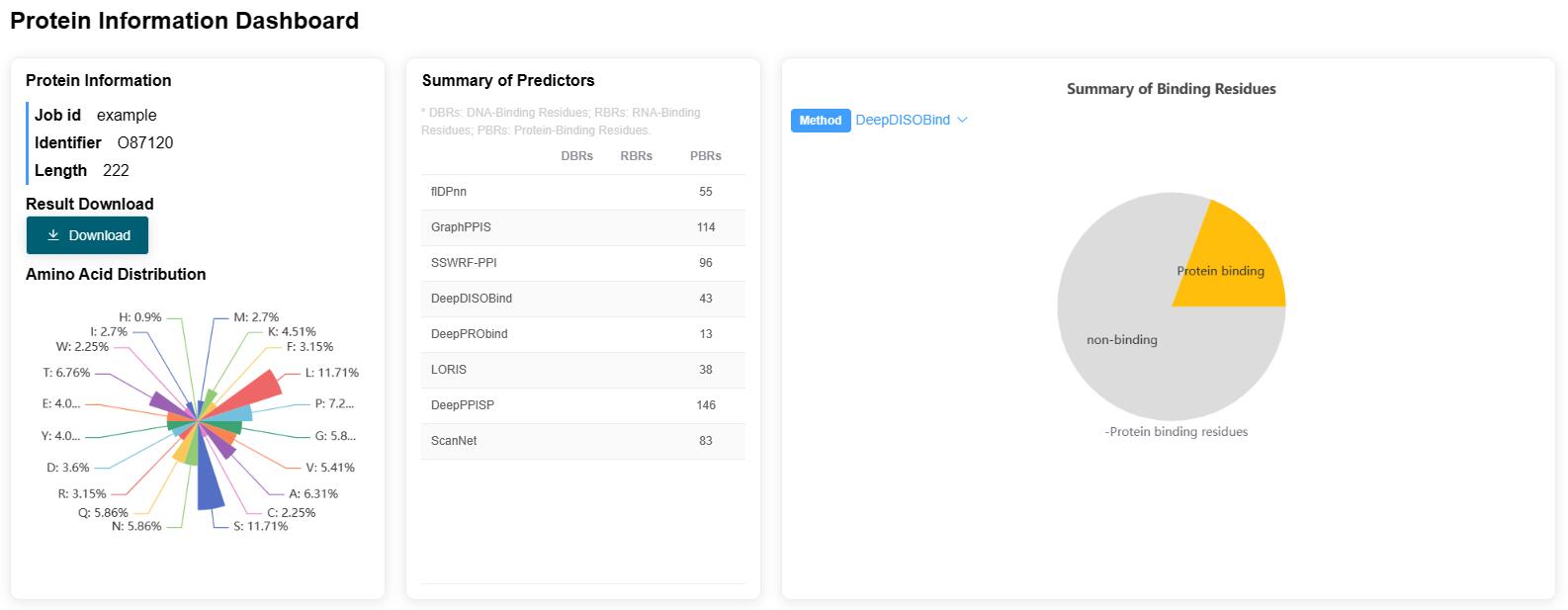

### Suppl. Figure S3

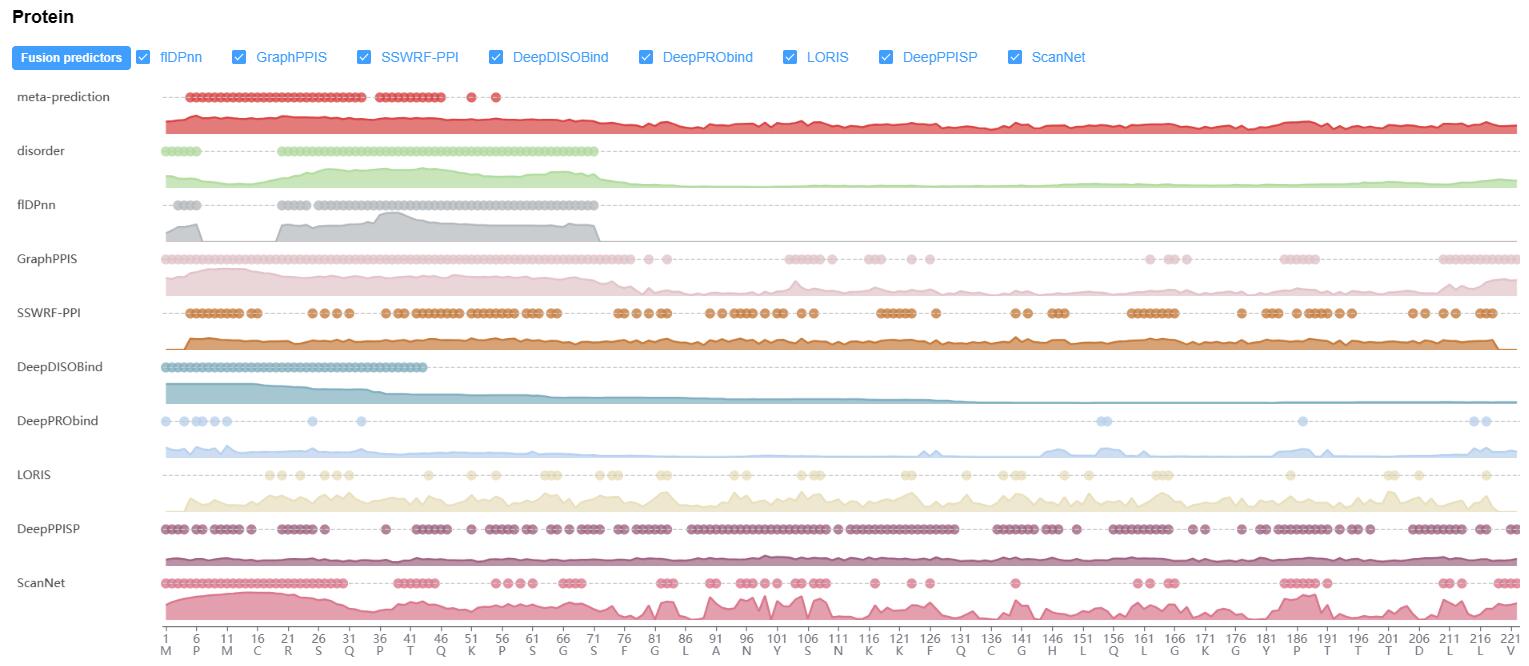

### Suppl. Figure S4

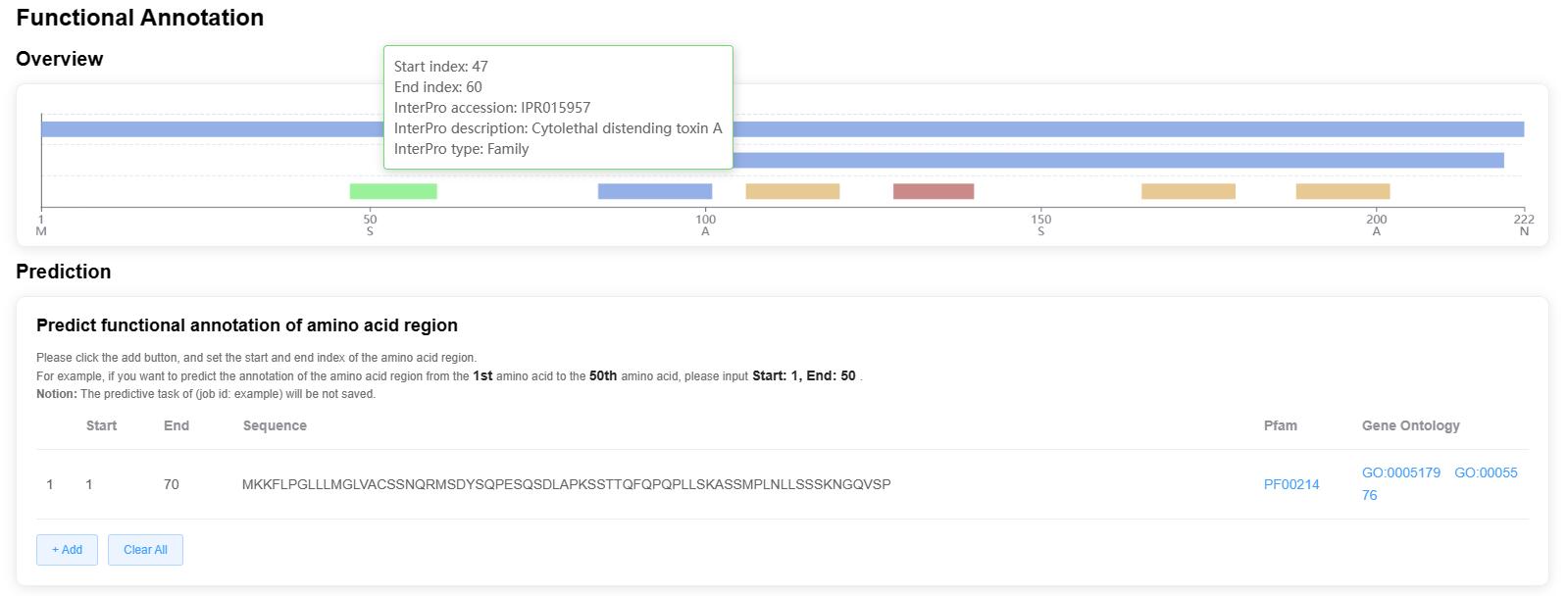

### Suppl. Figure S5

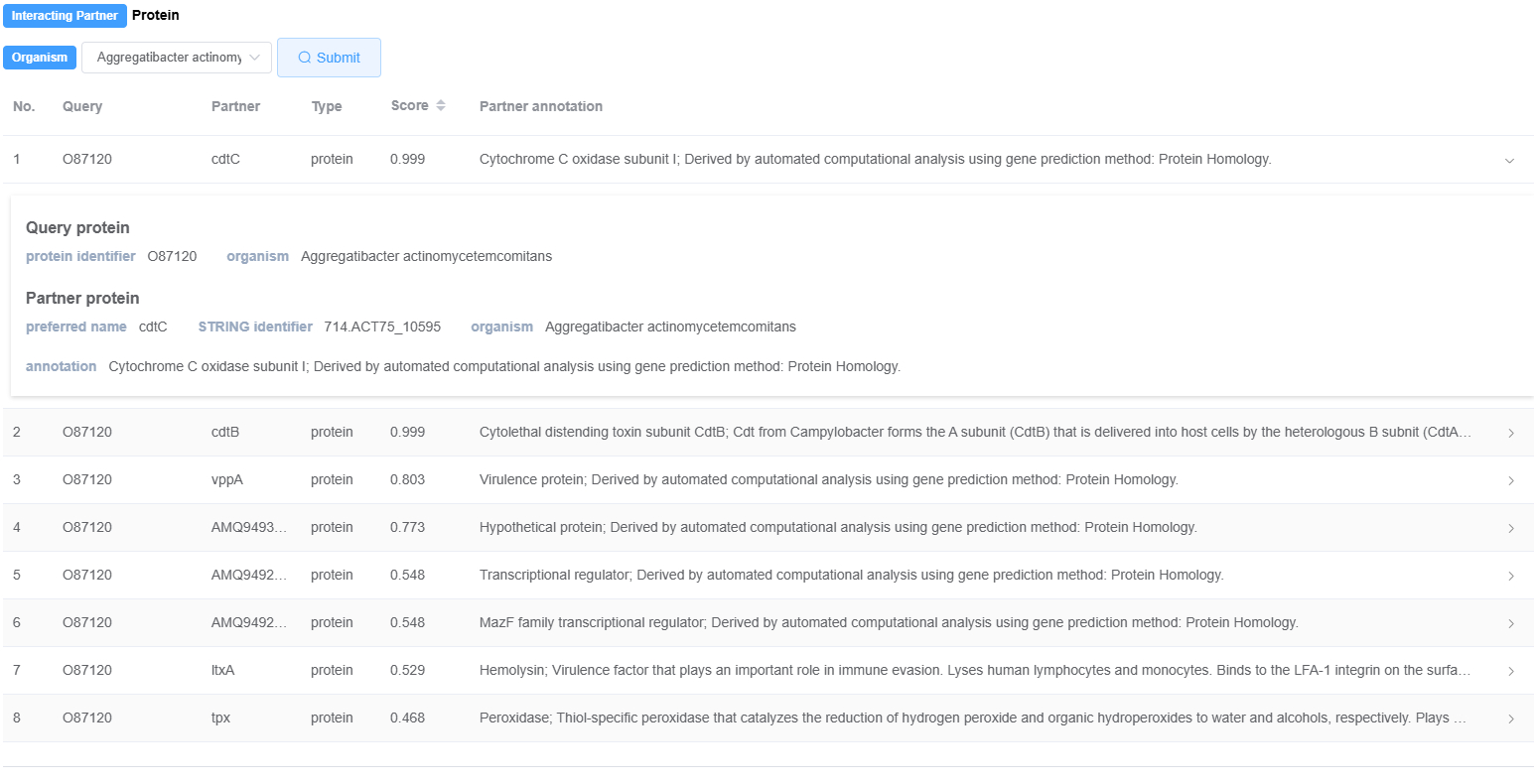

### Suppl. Figure S6

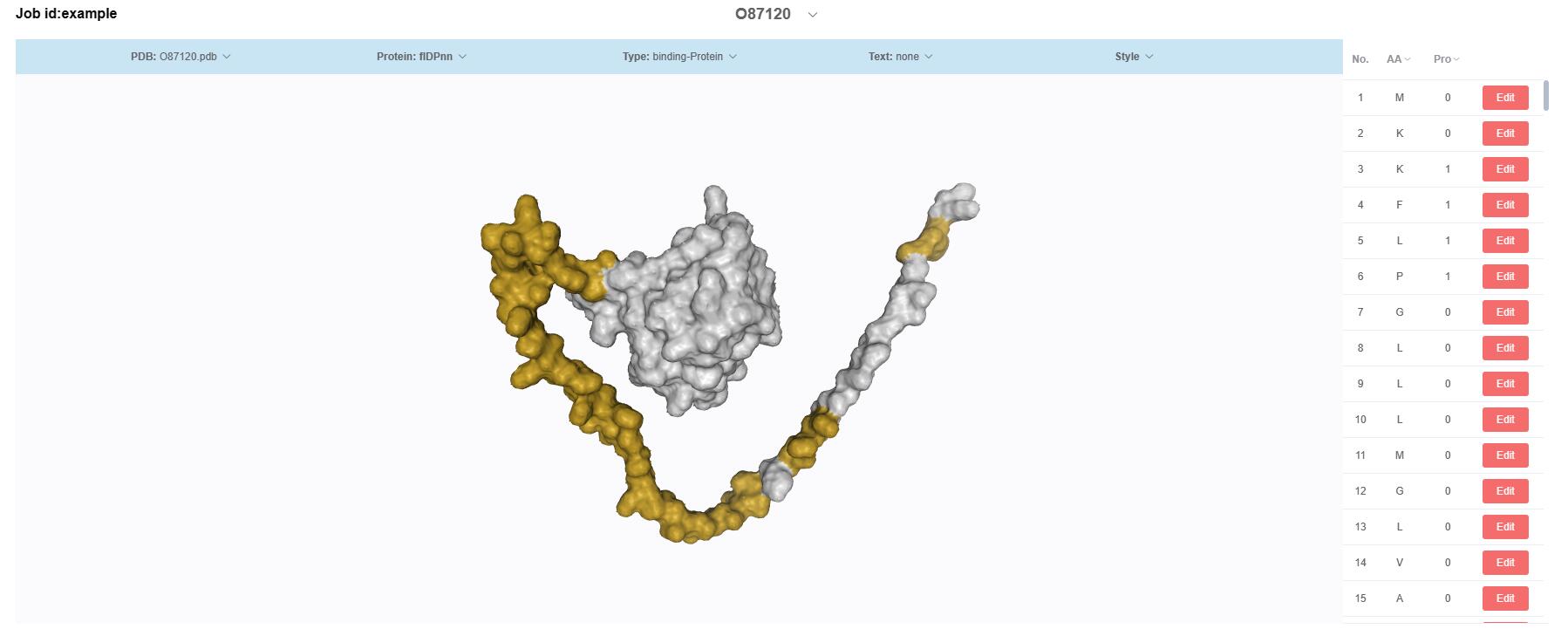
